# Escitalopram Administration, Neuroplastic Effects and Relearning: A Diffusion Tensor Imaging Study in Healthy Individuals

**DOI:** 10.1101/2021.04.25.441324

**Authors:** T Vanicek, MB Reed, J Unterholzner, M Klöbl, GM Godbersen, PA Handschuh, B Spurny, V Ritter, G Gryglewski, C Kraus, D Winkler, R Lanzenberger, R Seiger

## Abstract

**Background:** Neuroplastic processes are influenced by selective serotonergic reuptake inhibitors, while learning in conjunction with the administration of serotonergic agents alters white matter microstructure in humans. The goal of this double-blind, placebo-controlled imaging study was to investigate the influence of escitalopram on white matter plasticity during (re)learning.

**Methods:** Seventy-one healthy individuals (age = 25.6±5.0, 43 females) underwent 3 diffusion magnetic resonance imaging sessions: at baseline, after 3-weeks of associative learning (emotional/non-emotional content) and after relearning shuffled associations for an additional 3 weeks. During the relearning phase, subjects received daily escitalopram 10 mg or placebo orally. Statistical analysis was performed with statistical parametric mapping (SPM) and using sandwich estimator.

**Results:** A three-way and two-way rmANOVA was conducted to analyze the effects of escitalopram on AD, FA, MD and RD during the learning and relearning period. We found no significant three-way or two-way interactions for each DTI metrics (p_FDR_ > 0.05), thus neither after 3 nor after 6 weeks we found significant changes in white matter microstructure.

**Conclusion:** We examined neither an effect of escitalopram nor learning (or relearning) interventions on different DTI metrics. The duration and intensity of study interventions (i.e. administration of escitalopram and learning as the relearning task) might have been chosen insufficiently to induce detectable alterations. Previous studies examining the effects of SSRIs on white matter tracts in humans are underrepresented, but do mainly yield towards non-significant findings. The results implicate that escitalopram does not impact white matter microstructures in healthy subjects.

## Introduction

Antidepressants and in particular selective serotonin reuptake inhibitors (SSRIs) present a well-established and frequently applied treatment option for patients with mood and anxiety disorders (Bauer et al., 2013). Therapeutic agents modulating monoaminergic neurotransmission have been found to restore well-being with regard to different symptom complexes including depression, anxiety or obsessions and compulsions. According to the monoamine deficiency theory (Delgado, 2000), treatment-induced changes in mood following SSRI administration are physiologically related to a blockage of the serotonin transporter and subsequent up-regulated serotonin levels in the synaptic cleft that stimulates auto- and hetero-receptors located at serotonergic neurons and projection areas (Spies et al., 2015).

Throughout the last decade the influence of SSRIs on brain structure has been primarily investigated by exploiting T1-weighted magnetic resonance imaging (MRI). Most structural imaging investigations were conducted in patients with receiving SSRIs as a study and treatment intervention and results are partly heterogeneous. (Kraus et al., 2017). SSRI studies in healthy subjects and patients using MRI predominantly examine drug-induced changes in neuronal activation and functional connectivity (Dichter et al., 2015). Pharmacological studies on SSRI in healthy individuals and structural MRI are astonishingly scarce (Kraus et al., 2014; Shively et al., 2017). In longitudinal studies, cognitive or physical training and learning as well as administered SSRIs were found to impact brain morphology (Kristensen et al., 2018; Knorr et al., 2019). Also, SSRI intervention studies demonstrated treatment-associated changes in white matter integrity in different clinical samples with diverse methodological approaches (Yoo et al., 2007; Fan et al., 2012; Seiger et al., 2021).

Although SSRIs have been comprehensively studied on a cellular level in animal models and humans *in-vivo*, the mechanisms underlying their therapeutic potential in neuropsychiatric disorders are not well understood. Due to the partly insufficient and yet unexplained remission and response rates of approximately 40% to 60% following antidepressant treatment, respectively (Trivedi et al., 2006), the notion that SSRIs do not directly improve mood and thus cause depression to resolve, but instead enable susceptibility to change irrespective of direction has been increasingly studied and propagated (Chiarotti et al., 2017). SSRIs and fast-acting antidepressants as ketamine have been demonstrated to facilitate neuroplasticity (Alboni et al., 2017; Casarotto et al., 2021). According to this hypothesis, the brain is then suggested to be more prone to the quality and amplitude, implicating that the outcome of an SSRI treatment is dependent on encouraging or stressful influences one experiences (Branchi and Giuliani, 2021).

On the basis of imaging studies, convincing evidence points towards a relation between white matter integrity, plasticity and intellectual development of the central nervous system during child- and adulthood (Wang and Young, 2014; Bells et al., 2019). Apart from age-related changes of white matter throughout lifespan, which peak in the third and decline from the fifth decade onward (Dvorak et al., 2021), longitudinal neuroimaging studies revealed an effect of experience on white matter that can be traced with diffusion weighted imaging (Zatorre et al., 2012). Investigations in animals and humans show that white matter changes dynamically and dependent of the standardized study-experience *in vivo* following training (Keller and Just, 2009; Blumenfeld-Katzir et al., 2011). For instance, motor exercise learning as juggling increased fractional anisotropy (FA) in the right posterior intraparietal sulcus, an area associated with hand-eye co-ordination, and prevailed for at least 4 weeks following training (Scholz et al., 2009). The majority of white matter studies examined an increase in FA, and positive effects working memory on FA in parietal tracts (Takeuchi et al., 2010). In addition, perceptual learning such as texture discrimination learning or Braille reading training altered white matter, reflected in elevated FA (Yotsumoto et al., 2014; Debowska et al., 2016).

Within the last decade, diffusion tensor imaging (DTI) emerged as a reliable and frequently applied imaging approach to non-invasively investigate white matter microstructure. Although water diffusivity metrics are too coarse to comprehensively relate those findings to specific cellular mechanisms that are involved in white matter neuroplasticity, multimodal investigations in rodents have identified a significant overlap of molecular and cellular changes and white matter plasticity measured by diffusion MRI (Sampaio-Baptista et al., 2020). Several diffusion metrics have been found useful to describe differences between groups or over time (Smith et al., 2006). Among these metrics, FA provides information on the directionality of the diffusion process, hence, reflecting the grade of anisotropy and myelination. Axons allow water molecules to diffuse quickly along the direction of organization (axial diffusivity; AD), but constraining radial diffusivity (RD) meant to perpendicular travel along the direct axis. Mean diffusivity (MD) estimates the average water diffusion across all directions.

To reveal potential neuroplastic effects that SSRIs have on white matter during relearning, we performed a longitudinal placebo-controlled interventional learning study. Healthy subjects performed an associative learning task either with or without emotional content, where face pairs or pairs of Chinese characters and unrelated German nouns had to be memorized. After 21 days, subjects had to learn shuffled associations (i.e. relearning) within the same group for another 21 days. During the relearning period, subjects received either escitalopram 10 mg/day, or placebo. DTI models were estimated before and after associative learning and relearning periods.

## Methods

### Subjects and study design

Study participants were randomly assigned to one of four groups prior to study inclusion as follows: SSRI: 17 character-group, 16 faces-group; Placebo: 15 character-group, 23 faces-group. The design has been described in detail in previously published studies (Reed et al., 2021; Spurny et al., 2021). In brief, diffusion MRI scans were carried out before and after an associative learning period of 3 weeks as well as after a subsequent 3-week associative relearning under SSRI/placebo treatment. During the associative relearning period subjects received daily escitalopram 10 mg (Cipralex® Lundbeck A/S, provided by the Pharmaceutical Department of the Medical University of Vienna) orally or placebo. To ensure compliance, venous blood was drawn from the cubital vein to assess citalopram plasma through levels 1 week and 2 weeks into drug administration and before the third MRI. The therapeutic reference range for escitalopram it is 15-80 ng/mL (Hiemke et al., 2011).

The learning paradigm was divided into 2 groups (with or without emotional valence), where participants had to learn either face pairs or pairs of Chinese characters and unrelated German nouns. During the relearning phase the learning content was shuffled. During each phase, 200 image-pairs per group had to be learned or relearned with a daily subset of 52 image pairs. All faces were derived from the “10k Adult Faces Database” (Bainbridge et al., 2013).

### Data acquisition

The diffusion-weighted images (DWI) were acquired with a 3 Tesla Siemens MR scanner (MAGNETOM, Prisma, Siemens Medical, Erlangen, Germany) using a 64-channel head coil. 57 diffusion encoded images, with a b-value of 800 s/mm^2^ along with 13 non-diffusion (b=0) images were recorded (TR=9400 ms, TE=76 ms, slice thickness=1.6 mm, resolution=1.6⨯1.6 mm, matrix size=128⨯128⨯75, flip angle=90°, acquisition time=11:45 min). Subjects were instructed to keep their head as still as possible during the entire scan to minimize movement artifacts. Additionally, their heads were stabilized with foam cushions.

### Data processing

Data processing was conducted within the FMRIB software library (FSL, v. 5.0.11) (Smith et al., 2004) with the standard TBSS pipeline (Smith et al., 2006). All data outside of the brain were masked out (Smith, 2002), corrected for movements, geometric distortions and eddy currents as well as for outliers in the form of signal dropout (J. L. R. Andersson et al., 2016; J. L. R. Andersson and Sotiropoulos, 2016). The diffusion tensors were fitted using the dtifit command as well as the rotated b-vectors gained from the prior eddy current correction step. For the TBSS analysis, FA maps were eroded and brought into standard space using FNIRT (J. L. R. Andersson, Jenkinson, M., and Smith, S., 2007a; J. L. R. Andersson, Jenkinson, M., and Smith, S. M., 2007b). The data from each subject was then projected onto the FMRIB58_FA mean FA image and skeleton. The skeletonized images of all subjects were further used for voxel-wise statistics. In addition, the tbss_non_FA command was utilized to generate the skeletonized maps for the remaining parameters, AD, MD and RD.

### Statistical analysis

We applied an ANOVA as well as a Pearson’s chi-squared to test for similarities or differences between randomized groups. To reveal potential effects of escitalopram and (re)learning content have on both learning as well as relearning on DTI metrics, separate three-way repeated measures analyses of variance (rmANOVAs) for AD, FA, MD and RD were performed. We tested for interaction effects between substance (escitalopram vs placebo) x content (emotional vs. semantic learning and relearning) x time. Interaction effects were dropped in case of non-significance. To correct for different brain sizes and volumes the total intracranial volume was included as a covariate. The significance threshold was set at p⩽0.05 False discovery rate (FDR) was used to correct for all post-hoc comparisons. Statistical analysis was performed using the sandwich estimator (SwE) as implemented in statistical parametric mapping (SPM).

## Results

### Study population

Hundred and thirty-eight healthy subjects were included, but 54 subjects did not complete both the learning and relearning period (see Flow-diagram, Table 1). Data from 71 healthy subjects (age = 25.6 ± 5.0) measured at three time points in a randomized, double blind, placebo-controlled manner were included in this DTI analysis. The four groups did not significantly differ in terms of age and gender, as shown by the ANOVA: F (3,67) = 0.025, p=0.995 and the Pearson’s chi-squared test: X-squared = 2.24, df = 3, p-value = 0.524. Detailed demographical results can be found in Table 2.

**Table 1:**
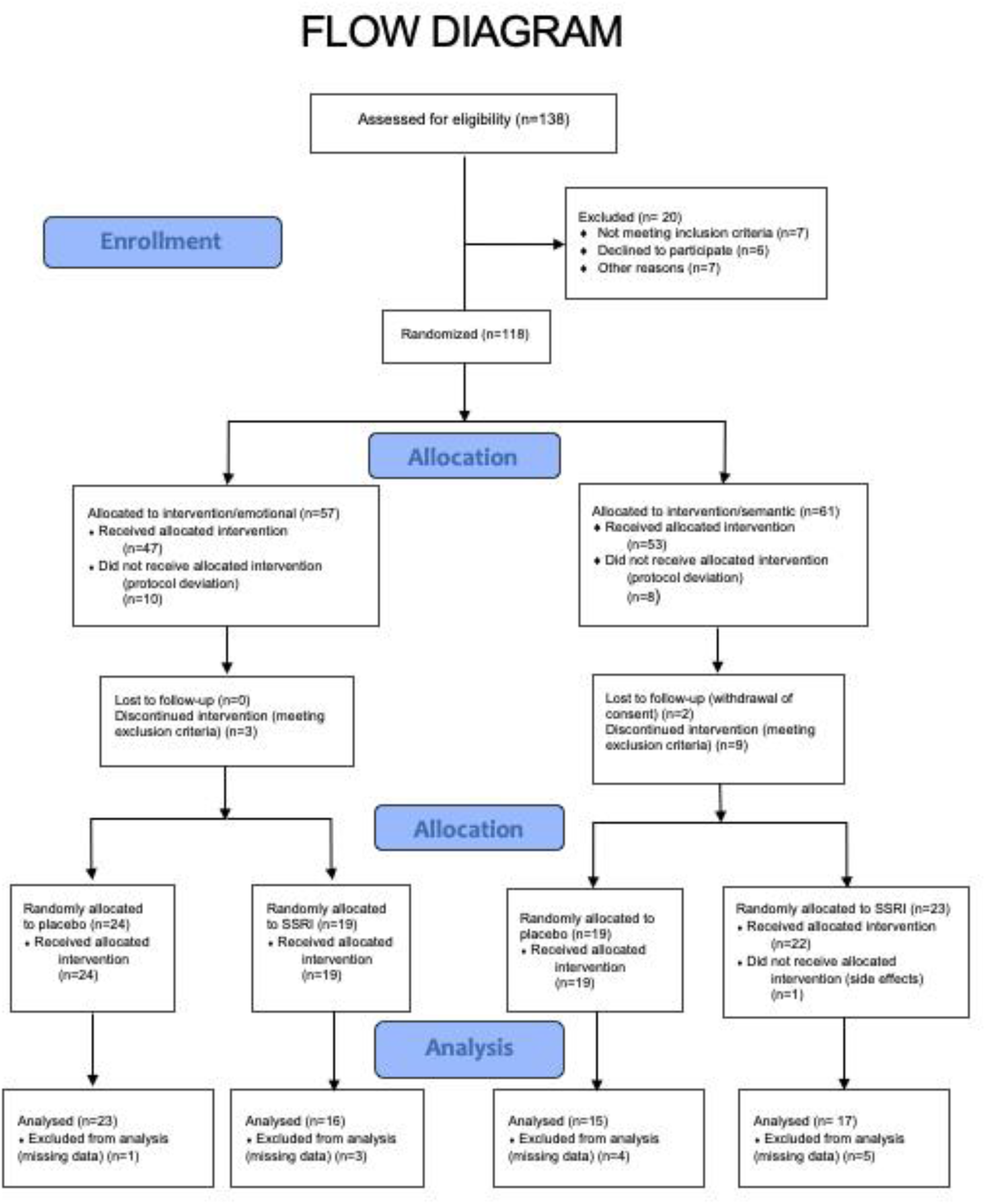
Flow-Diagram.

**Table 2:**
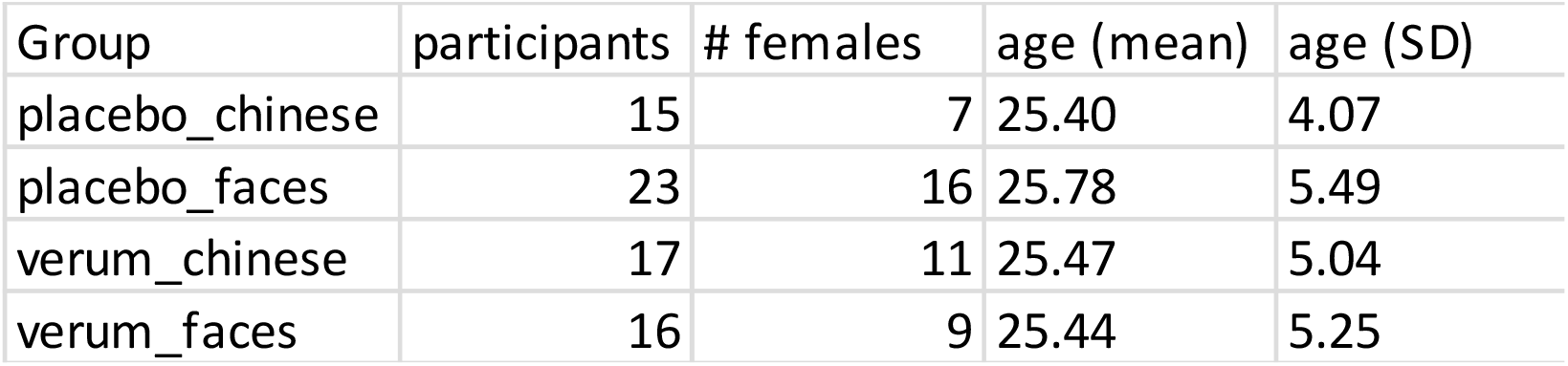
Demographical information for subjects included in DTI analysis.

### Analysis of DTI metrics over time

To test if escitalopram (vs. placebo) effects AD, FA, MD and RD during the learning and relearning period and if learning content (emotional vs. semantic) is relevant, a three-way and two-way rmANOVA were conducted first to analyze interaction effects. We observed no significant three-way interactions for each DTI metrics (p_FDR_ > 0.05). Further, we also found no interaction between learning as well as relearning and time and no interaction between substance and time.

## Discussion

Given preclinical reports on the neuroplastic effects of SSRI and its relevancy for neuropsychiatric disorders, we aimed to investigate the influence of escitalopram on white matter microstructure *in vivo* during relearning. We examined no influence of escitalopram on DTI metrics during the relearning period. Further, neither learning (2^nd^ MRI at day 21) nor relearning (3^rd^ MRI at day 42) lead to significant differences in AD, FA, MD and RD. Here, learning content (emotional or semantic content) also had no influence on white matter.

Work by our group demonstrated immediate changes in diffusivity parameters following intravenous SSRI-administration. In a PET/MRI intervention study, patients with depression and healthy controls received citalopram 8 mg intravenous, which was described to sufficiently occupy the serotonin transporter (Gryglewski et al., 2019). Citalopram led to attenuated MD, AD and RD in the anterior corona radiata, corpus callosum, external capsule and frontal blade in patients and controls, without affecting FA or increasing any DTI metric (Seiger et al., 2021). Also, short and intensive learning has been found to affect white matter microstructure (Hofstetter et al., 2013), suggesting that cellular rearrangement in this compartment occurs rapidly (Sagi et al., 2012).

In comparison to these short-time effects of SSRI, here, we administered escitalopram over a period of 3 weeks and depicted no effects on white matter microstructure. In a DTI study in patients with obsessive-compulsive disorder, decreased MD and RD in midbrain regions and striatum were observed following 12 weeks of SSRI treatment, while changes in FA were not detected (Fan et al., 2012). These findings link SSRI to diminished FA and AD, indicating a reduction in longitudinal water diffusion. Also, disorder-specific symptoms decreased significantly following 12 weeks of citalopram treatment in patients with obsessive-compulsive disorder in comparison to controls, while FA was higher before treatment initiation and diminished over time in the corpus callosum, internal capsule and adjacent structures to the right caudate. However, the findings were not corrected for multiple comparisons (Yoo et al., 2007). Effects of SSRI on diffusivity parameters in patients with depression were found divergent in two studies. One study found unspecific white matter volume alterations across different areas after 12 to 16 weeks of SSRI-treatment in 17 patients with depression, though, only if significant levels were chosen less conservative (Zeng et al., 2012). In a multicenter study 200 patients with depression were measured 3 times across 8 weeks. Interestingly, Davis et al. found changes at baseline in comparison to control subjects, but no longitudinal white matter changes across the 8-week timespan (Davis et al., 2019). Our study findings appear to reflect these reports, since SSRI interventions have been mainly found to not propagate changes of DTI metrics, in particular when analyses are performed with conservative measures. Then, the timespan of 3 weeks of escitalopram administration was distinctively shorter than in other white matter studies, where certain changes over time were demonstrated. Nevertheless, this duration was found reasonable for SSRIs to promote neuroplastic effects and unfold efficacy in clinical settings (Bauer et al., 2013; Alboni et al., 2017).

For long, human behavior and perception was understood as an effector of neuronal activity, with the synapse as the primary location for cell communication and thus neuroplasticity, while white matter was reduced to a (in-)homogeneous structure serving as a medium for proper transfer of information. Intervention studies led to the conception that the plasticity phenomena not only occurs in grey matter, where environmental stimuli are known to shape synaptic organization (Draganski et al., 2004), but are relevant for the reorganization of white matter likewise (Sampaio-Baptista and Johansen-Berg, 2017). Activity-dependent changes of myelination architecture are found to affect neuronal networks (Hartline and Colman, 2007; Ford et al., 2015). Non-pharmacological interventions as motor skill learning or cognitive training have been found to be associated with DTI-changes in healthy subjects and patients with mental disorders. These effects were demonstrated to be dependent on time or training load, since interventions with an interval less than 8 weeks did not generate pre-post differences in white matter (Kristensen et al., 2018). Here, we examined no effects of learning or relearning intervention on DTI metrics, indifferent if learning groups with emotional or semantic content were compared or learning groups were pooled together. In the same study sample, metabolic and functional changes have demonstrated after relearning and escitalopram administration. In a task-based fMRI analysis, neuronal activation was found to be decreased in the insula after relearning in subjects that were administered escitalopram (Reed et al., 2021). Further, we found a relationship between escitalopram and relearning when comparing glutamate levels in the hippocampus and thalamus before and after associative relearning (Spurny et al., 2021). These findings underline the effects of escitalopram and learning on various brain measures and we expected learning induced white matter changes. Nevertheless, it is probable that learning intensity and duration were designed to sparse to propagate adaptations in the white matter compartment.

Within the group that received escitalopram during reversal learning, half of the subjects developed citalopram blood levels beneath the therapeutic range. In clinical studies, patients receiving SSRIs have generally not been found to significantly benefit from dose escalation strategies, since response rates are found similar for therapeutic dose and high-dose treatment, but less tolerability and acceptability occurred in treatment with higher doses (Adli et al., 2005; Furukawa et al., 2019). Already very low doses of citalopram, the racemic precursor substance of escitalopram, lead to a serotonin transporter occupancy of 50 %, while 80 % occupancy was depicted at therapeutic citalopram dosing, that is equivalent to the applied dose of escitalopram (i.e. 10 mg daily) in our study (Meyer et al., 2004; Shapiro, 2018). However, antidepressants are found to induce neuronal plasticity in dependence of drug dose in clinical samples (Chiarotti et al., 2017). Taken together, therapeutic doses of escitalopram 10 mg should have sufficiently stimulated serotonergic neurotransmission and prevented from higher drop-out rates, though mediocre citalopram blood levels across our study individuals could have affected study outcomes.

We aimed to reveal a relationship between SSRI treatment, neuroplastic effects on white matter properties and reversal learning in humans *in vivo*. Results show neither an effect of escitalopram nor learning or relearning interventions on DTI metrics. The duration and intensity of study interventions, i.e. administration of escitalopram and learning as the relearning task, might have been chosen insufficiently to induce detectable effects.

## Acknowledgements

We wish to thank the team members as well as the medical students of the Neuroimaging Lab (NIL), headed by Prof. Lanzenberger, the corresponding author. Also, we thank T. Stimpfl for SSRI plasma level analyses.

## Ethics approval

This study is part of a larger project that has been approved by the ethics committee of the Medical University of Vienna (EK Nr.: 1739/2016) and performed in accordance with the Declaration of Helsinki (1964). The project is registered at clinicaltrials.gov with the identifier NCT02753738.

## Availability of data and material

The full data can be made available upon reasonable request to the corresponding author.

